# Identification of a novel deFADding activity in 5’ to 3’ exoribonucleases

**DOI:** 10.1101/2022.05.10.491372

**Authors:** Sunny Sharma, Jun Yang, Selom K. Doamekpor, Ewa Grudizen-Nogalska, Liang Tong, Megerditch Kiledjian

**Affiliations:** Department of Cell Biology and Neurosciences, Rutgers, University, Piscataway, New Jersey, 08854, USA; Department Biological Sciences, Columbia University, New York, NY 10027, USA

## Abstract

Identification of metabolite caps including FAD on the 5’ end of RNA has uncovered a previously unforeseen intersection between cellular metabolism and gene expression. To understand the function of FAD caps in cellular physiology, we characterised the proteins interacting with FAD caps in budding yeast. Here we demonstrate that highly conserved 5’-3’ exoribonucleases, Xrn1 and Rat1, physically interact with the RNA 5’ FAD cap and both possess FAD cap decapping (deFADding) activity and subsequently degrade the resulting RNA. Xrn1 deFADding activity was also evident in human cells indicating its evolutionary conservation. Furthermore, we report that the recently identified bacterial 5’-3’ exoribonuclease RNase AM also possesses deFADding activity that can degrade FAD-capped RNAs *in vitro* and in *E. coli* cells. To gain a molecular understanding of the deFADding reaction, an RNase AM crystal structure with three manganese ions coordinated by a sulfate molecule and the active site amino acids was generated that provided details underlying hydrolysis of the FAD cap. Our findings reveal a general propensity for 5’-3’ exoribonucleases to hydrolyse and degrade RNAs with 5’ end noncanonical caps in addition to their well characterized 5’ monophosphate RNA substrates indicating an evolutionarily conserved intrinsic property of 5’-3’ exoribonucleases.

## INTRODUCTION

Regulation of gene expression in consonance with metabolic state is critical for cell survival. This coordination between metabolism and gene expression is driven by several key metabolites, particularly adenine-containing nucleotide metabolites such as Nicotinamide Adenine Dinucleotide (NAD), Flavin Adenine Dinucleotide (FAD), Coenzyme A, and S-adenosyl methionine (SAM) (1). These metabolites generally function as cofactors or co-substrates for many enzymes that modify chromatin and play key tasks in both activation and repression of gene expression (2).

Recent identification of NAD as an RNA 5’ end cap in bacteria (3), yeast (4), plants (5,6), and mammals (7) has uncovered a unique metabolic network where a metabolite can directly influence gene expression. Both structural and biochemical analyses in prokaryotes have revealed that RNA polymerases can use NAD in place of ATP at the 5’ end during transcription initiation (8). In contrast, a NAD cap on post-transcriptionally processed intronic small nucleolar RNAs (snoRNAs) in mammalian cells has highlighted an alternative mechanism of a post-transcriptional NAD capping (7). Recent studies in model organisms representing all three kingdoms of life have led to the identification of different NAD cap decapping (deNADding) enzymes that has aided in understanding the biological function of NAD caps (3,5–7,9–11). In prokaryotes, specifically in *E. coli*, the NAD cap protects RNA from 5’ end degradation (3), whereas in mammals, the NAD cap serves as a mark to degrade the RNA at least in part by the DXO 5’-3’ exoribonuclease (7). More recently, characterization of NAD cap binding proteins in yeast led to the identification of the highly conserved Xrn1 and Rat1 5’-3’ exoribonucleases as novel deNADding enzymes and revealed a central role of NAD capped RNAs in mitochondrial function (11).

In addition to NAD caps, chemical analysis of the RNA 5’-cap epitranscriptome using LC-MS/MS based CapQuant (12) from different organisms revealed that mRNAs also contain other metabolite caps like FAD, dephosphoCoenzyme (dpCoA), Uridine diphosphate glucose (UDP-Glc), and Uridine diphosphate *N*-acetylglucosamine (UDP-GlcNAc). Nevertheless, the function of these metabolite caps in RNA metabolism remained elusive. It is important to note that identification of the enzymatic machinery involved in addition and removal of these metabolite caps is imperative to understanding their biological function. Our initial screening of putative metabolite cap decapping enzymes for FAD cap (deFADding), led to the identification of Rai1 along with its mammalian homolog DXO (13), and two NUDIX proteins-Nudt2 and Nudt16 as deFADding enzymes (10).

In the present study, we performed FAD cap RNA affinity purification (FcRAP) and identified novel FAD cap associated proteins in budding yeast. Interestingly, in addition to their role in hydrolyzing NAD-capped RNA (11), Xrn1 and Rat1 are potent deFADding enzymes. Furthermore, a recently reported 5’-3’ exoribonuclease, RNase AM (yciV) (14,15) in *E. coli* is also capable of deFADding and hydrolyzing FAD capped RNA and its structure reveals the organization of its active site. Our studies suggest a potential commonality of 5’-3’ exoribonucleases towards FAD-capped RNA hydrolysis and decay in both prokaryotes and eukaryotes.

## MATERIAL AND METHODS

### FAD cap RNA Affinity Purification (FcRAP)

The *in vitro* transcribed FAD and m^7^G capped RNAs (^~^400 pmol) were immobilized onto Dynabeads™ MyOne™ Streptavidin T1 (Invitrogen™)). Yeast cells (Supplementary Table S3) were grown and processed as described recently (11). Briefly, 5 mg of protein lysate was added to the preequilibrated (in 1X lysis buffer) FAD- or m^7^G-capped RNA linked Dynabeads in 2 mL microcentrifuge tubes. These tubes were next incubated at 4°C on an end-to-end rocker for 90 minutes. After extensive washing (at least 5 times with 1X lysis buffer) the associated proteins were eluted with 2X Laemmli buffer (Bio-Rad Laboratories). The samples were denatured at 85°C for 5 minutes and were next run on Novex™ WedgeWell™ 4 to 20 %, Tris-Glycine Protein Gel (Thermo Fisher Scientific). The protein gel was stained with SYPRO Ruby (Bio-Rad Laboratories) or with silver staining as described before (16), and specific bands were sliced and processed for mass spectrometry-based identification. In addition to the individual sliced bands, we also examined the total eluate using mass spectrometry. Complete list of identified proteins is provided in the Supplementary Table S1.

Mass spectrometry was carried out at Biological Mass Spectrometry Facility of Robert Wood Johnson Medical School and Rutgers, The State University of New Jersey and as described previously (11).

### Electrophoretic Mobility Shift Assay

Recombinant *K. lactis* Xrn1 (2.5, 5 and 7.5 pmol) was mixed with a fixed concentration (5 pmol) of ^32^P uniformly-labeled NAD- or FAD- or pA-in vitro transcribed RNA in 1x RBB buffer (10 mM Tris-HCl pH7.5, 150 mM KCl, 1.5 mM MgCl_2_ and 0.5 mM DTT) and incubated on ice for 30 minutes. The samples were resolved on to a 6% acrylamide gel using 0.5x TBE at 120V for 2-3 hrs. The dried gel was exposed to phosphor screen overnight and visualized with an Amersham Typhoon RGB Biomolecular Imager (GE Healthcare Life Sciences).

### Plasmid construction and site-directed mutagenesis

All proteins used in the present study for *in vitro* assays were purified as described previously (17). For RNase AM, complete ORF was cloned into a modified pET28a vector, in-frame with an N-terminal 6xHis-tagged yeast SMT3 SUMO gene using In-Fusion^®^ Snap Assembly (TaKaRa Bio). For the construction of different point mutants, site directed mutagenesis were performed using SPRINP (18) as described previously using the oligonucleotides listed in the Supplementary Table S2.

### Protein Expression and Purification

All recombinant proteins – Kl Xrn1, Hs Xrn1 and RNAse AM along with the mutants were expressed and purified using Nickel-NTA affinity purification (Supplementary Table S3). 10 mL LB cultures containing 50 μg/mL kanamycin were inoculated with single colonies and grown overnight. For purification, 1 L LB culture was gown at 37°C to an OD_600_ of 0.5. Protein expression was induced with 0.25 mM isopropyl D-thiogalactopyranoside (IPTG) and cells grown at 18°C on a shaker for ~18 h. Cells were collected by centrifugation at 5000 g for 15 minutes, washed with PBS. For protein purification, cell pellets were resuspended in ~60 mL of buffer (25 mM HEPES (pH 7.5), 250 mM KCl, 10% glycerol, and 100 μM MnCl_2_) containing 0.4 mg/mL protease inhibitor phenylmethanesulfonylfluoride fluoride (PMSF). Cells were lysed by sonication, and the insoluble cell debris was separated by centrifugation. The Nickel-NTA affinity purification was performed as described previously (17).

### *In vitro* transcription of NAD-, FAD-capped RNAs

NAD, FAD, and m^7^G cap containing RNAs were synthesized by *in vitro* transcription from a synthetic double stranded DNA template ϕ2.5-NAD/FAD-40 containing the T7 ϕ2.5 promoter and a single adenosine within the transcript contained at the transcription start site (Supplementary Table S2). For m^7^G-capped RNA, m^7^GpppA RNA Cap Structure Analog (New England Biolabs) was included in the transcription reaction. *In vitro* transcription was carried out at 37°C overnight, using HiScribe™ T7 High yield RNA Synthesis kit (New England Biolabs). To generate ^32^P uniformly labeled RNA, the transcription reactions were performed in the presence of [α-^32^P] GTP.

### RNA *in vitro* deFADding assay

The [α-^32^P] GTP uniformly labeled FAD-capped RNAs were incubated with the specified amounts of recombinant protein in NEB buffer 3 (100 mM NaCl, 50 mM Tris-HCl, 10 mM MgCl_2_, pH 7.9). For RNase AM, 4 mM MnCl_2_ was used instead of MgCl_2_. Reactions were stopped by heating at 85°C for 2 minutes. *In vitro* 5’end RNA decay was carried out as described (11), cell extract corresponding to 2 μg of cellular protein from WT or *xrn1Δ* strains were incubated with [α-^32^P] GTP uniformly labeled FAD and m^7^G-capped RNA immobilized onto Dynabeads™ MyOne™ Streptavidin T1 (Invitrogen™)) in NEB buffer 3.

### FAD-cap detection and quantitation (FAD-capQ)

Bacterial and yeast total RNA were extracted using hot phenol (19), whereas RNA from human cells were isolated using Trizol (Invitrogen™)). The strains used in the present study are listed in Supplementary Table S3. To remove residual free FAD, purified RNAs were dissolved in 10 mM Tris-HCl (pH 7.5) containing 2 M urea. RNA samples were incubated 2 min at 65°C and immediately precipitated with isopropanol in the presence of 2 M ammonium acetate. FAD-capQ was carried out as previously described (13). For human cells, 30 μg of small RNAs (<200 nts) isolated using Monarch (NEB) were used. For bacteria and yeast, 50 μg of total RNA was used for the FAD-capQ analysis.

To demonstrate the release of intact FAD upon Xrn1 or RNAse AM mediated deFADding of FAD capped RNAs, FAD-capQ assay using different amounts of *in vitro* transcribed FAD capped RNA was used. For the Xrn1 hydrolysis reactions, 100 mM of the WT or E178Q catalytic-inactive mutant Xrn1 was used. For RNase AM, 150 nM of WT or catalyically-inative D20A mutant was used.

### Structure determination

An expression construct for full-length *E. coli* RNase AM was used to transform *E. coli* BL21 (DE3) Rosetta cells. His-tagged RNase AM was produced and purified like previously reported (14). Prior to induction with 0.25 mM IPTG, the iron-specific chelator 2,2’-bipyridyl was added to a final concentration of 0.15 mM for 30 minutes followed by the addition of MnSO_4_ at 1 mM. After cell disruption by sonication and centrifugation, the lysate (in a buffer containing 250 mM NaCl, 20 mM Tris (pH 7.5), 20 mM, imidazole, 5 mM β-mercaptoethanol (BME), 10% (v/v) glycerol, 0.1 mM MnSO_4_, and 1 mM phenylmethanesulfonylfluoride (PMSF)) was incubated with 2 ml of Ni-NTA superflow resin (Qiagen) with nutation for 1 h. RNase AM was eluted with 5 ml of buffer containing 250 mM NaCl, 20 mM Tris (pH 7.5), 250 mM imidazole, 5 mM BME, 10% (v/v) glycerol and 0.1 mM MnSO_4_ and further purified with gel filtration (Superdex 200, Cytiva Life Sciences) chromatography (in buffer containing 200 mM NaCl, 20 mM Tris (pH 7.5) and 2 mM DTT).

Attempts to crystallize RNase AM alone or supplemented with FAD did not yield diffraction quality crystals. Good-quality RNase AM crystals were obtained in the presence of dsRNA (from the CDE of IL6) using the sitting-drop vapor diffusion method at 20°C with a reservoir solution containing 20% (w/v) PEG 8000 and 0.1 M Hepes pH 7.0. RNase AM crystals were cryo-protected with 20% (w/v) PEG 8000 and 25% (v/v) ethylene glycol before being flash frozen in liquid nitrogen for diffraction analysis and data collection at 100 K. X-ray diffraction data were collected at the Advanced Photon Source (APS) beamline 24-ID-C. The diffraction images were processed and scaled with standard parameters using the XDS program (20). The crystal belongs to space group *P2_1_*.

Molecular replacement (MR) using full-length amidohydrolase from *Chromobacterium violaceum* (PDB: 2YB1) (21) did not yield a molecular replacement solution, with the program Phaser (22). A correct MR solution could be obtained only with the PHP domain. No additional density for the insertion domain of RNase AM was observed, suggesting that it is disordered. The structure refinement was performed using PHENIX (23) and manual rebuilding of the atomic model was carried out with the Coot program (24). The final model contains residues 9-106 and 177 to 284. Density for the 3 metal ions was assigned as manganese with a sulfate molecule in the active site and no RNA density was observed. The crystallographic information is summarized in Table 1.

**Table 1.** Summary of crystallographic information.

Summary of crystallographic information
| Structure | RNase AM in complex with Mn and Sulfate |
| --- | --- |
| **Data Collection** |  |
| Space group | $P2_1$ |
| Cell dimensions |  |
| $a, b, c$ (Å) | 40.3, 74.4, 40.1 |
| $\alpha, \beta, \gamma$ (°) | 90, 119.0, 90 |
| Resolution (Å) | 35.24-1.70 (1.75-1.70) |
| $R_{\text{merge}}$ (%) | 13.7 (74.4) |
| $CC_{1/2}$ (%) | 98.7 (78.1) |
| $I/\sigma I$ | 10.0 (3.9) |
| Completeness (%) | 97.7 (94.3) |
| No. of reflections | 22722 (2180) |
| Redundancy | 5.0 (4.9) |
| **Refinement** |  |
| Resolution (Å) | 35.2-1.70 (1.75-1.70) |
| $R_{\text{work}}/R_{\text{free}}$ (%) | 18.2 (35.4) / 21.3 (41.8) |
| Number of atoms | 1768 |
| Protein | 1631 |
| Ligand/Ion | 8 |
| Water | 129 |
| B-factors (Å<sup>2</sup>) | 23.3 |
| Protein | 22.7 |
| Ligand | 18.7 |
| Water | 31.1 |
| r.m.s.d. |  |
| Bond lengths (Å) | 0.007 |
| Bond angles (°) | 1.00 |
| Ramachandran plot statistics |  |
| Favored | 97.0 |
| Allowed | 3.0 |
| Outliers | 0.0 |

## RESULTS

### Characterization of FAD cap binding proteins in budding yeast

We recently employed NAD cap RNA affinity purification (NcRAP) to characterize NAD cap binding proteins to study their biological roles (11). Similarly, to begin understanding the functional role of FAD caps, we performed 5’ FAD cap RNA Affinity Purification (FcRAP). Here, an *in vitro* transcribed FAD-capped RNA containing a 3’-biotin was used as bait to capture and identify proteins associated with the FAD cap from *S. cerevisiae* lysate as illustrated in Figure 1A. Two identical RNAs that differed only at their 5’ end cap (m^7^G or FAD cap) were used to account for common RNA-binding proteins. Affinity purified proteins associated with the different RNAs were identified by mass spectrometry and the m^7^G-capped RNA was used as the control. Mass spectrometry identification of proteins selectively bound to the FAD-cap (Supplementary Data 1) revealed the most prominent FAD cap associated proteins to be the 5’-3’ exoribonucleases Xrn1 and Rat1 as well as the Rat1 associated deFADding protein, Rai1 (13,25) (Figure1B). FcRAP was also carried out with protein extract derived from an Xrn1 deletion (*xrn1Δ*) strain which still retains the association of Rat1 and Rai1 but as expected is devoid of Xrn1 association (Figure 1B). Lastly, to control for the association of Rat1 to the FAD cap through its interacting protein Rai1, a similar analysis was carried out with extract devoid of Rai1 (*rai1Δ*) where Rat1 is still detected with the FAD-capped RNA (Supplementary Figure S1). These findings demonstrate all three proteins have the capacity to associate with the FAD cap. Interestingly, these three proteins were also recently shown to selectively associate with and hydrolyze the NAD cap (11).

**Figure 1.**
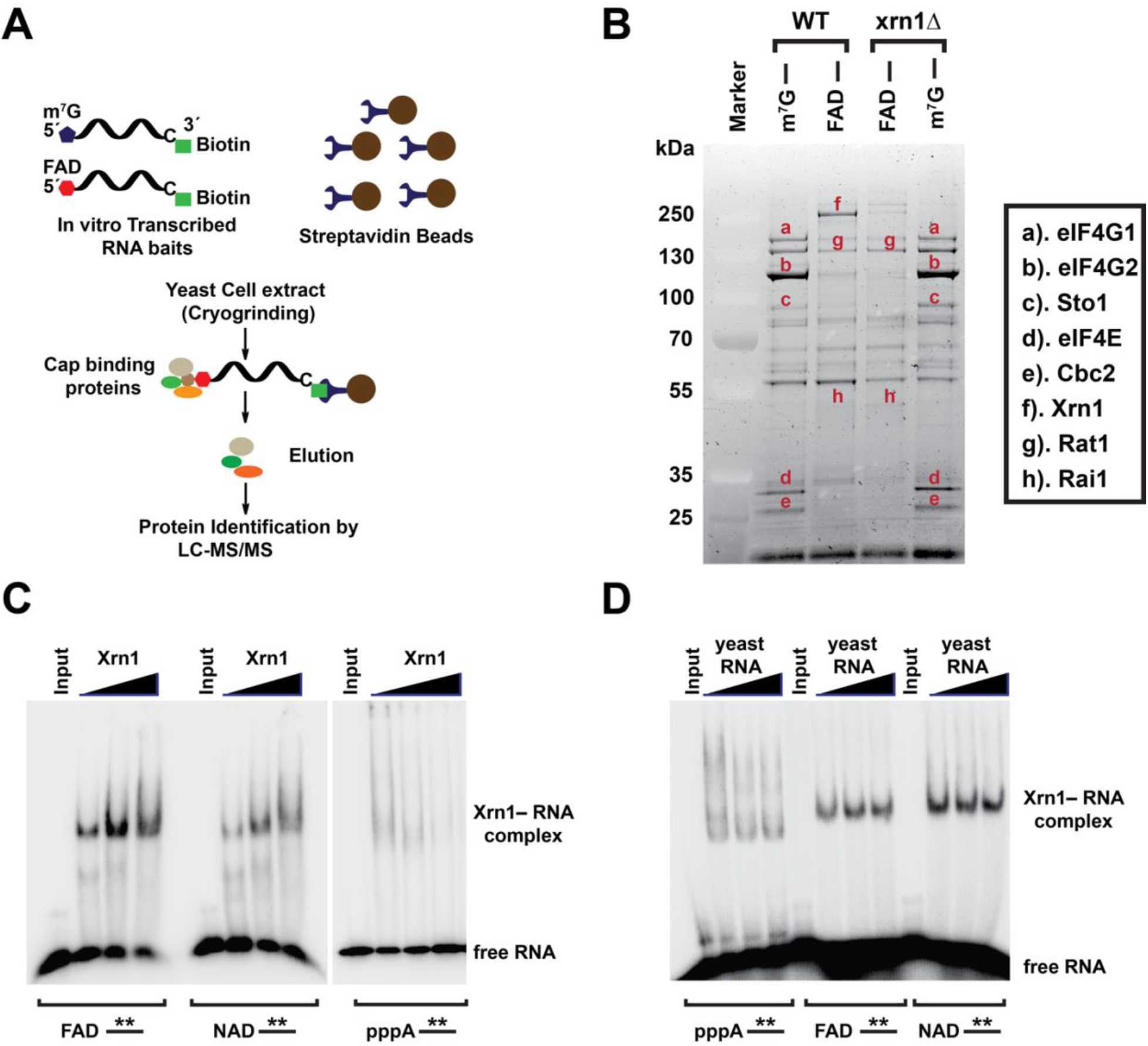
Identification of FAD cap binding proteins in budding yeast. (A) Schematic illustration of the FAD cap -RNA Affinity Purification (FcRAP). (B) Eluates from FcRAP were loaded on to a 4%-12% Bis-Tris gel and stained with SYPRO Ruby. The m^7^G cap affinity purification was used as a control. All protein bands labeled on the gel were excised from the gel and identified using mass spectrometry. (C) Electrophoretic mobility shift assay (EMSA) using increasing concentration of Xrn1 (2.5 pmol, 5 pmol and 7.5 pmol) and a fixed concentration (5 pmol) of uniformly labeled *in vitro* transcribed FAD-capped or NAD-capped or triphosphate RNA (pppA–). (D) EMSA using equimolar concentration (5pmol) of Xrn1 and uniformly labeled *in vitro* transcribed FAD-capped or NAD-capped or pppA–RNAs in the presence of total yeast RNA as a nonspecific competitor.

The interaction of Xrn1 with the FAD cap was further explored using electrophoretic mobility shift assays (EMSAs). Equimolar concentration of Xrn1 and uniformly labeled *in vitro* transcribed FAD and NAD capped RNAs along with triphosphate RNA were mixed and incubated on ice for 30 minutes. Xrn1 can form a stable complex with both FAD and NAD-capped RNA (Figure 1C) that is minimally competed with nonspecific competitor total yeast RNA (Figure 1D). Use of an identical control RNA containing a 5’ triphosphate demonstrated nonspecific associations that were more readily competed by the competitor RNA. These findings reveal Xrn1 is capable of selectively binding to both NAD and FAD caps under the assay conditions employed.

### Xrn1 and Rat1 hydrolyze FAD capped RNAs *in vitro*

The capacity for Xrn1 and Rat1 to hydrolyze a FAD cap (Figure 2A) was next tested. This was predicated on our recent demonstration that both enzymes can hydrolyze NAD-capped RNA (11). Uniformly ^32^P-labeled FAD-capped RNA was degraded by wild-type *Kluyveromyces lactis* Xrn1 and *Schizosaccharomyces pombe* Rat1, but not their corresponding catalytically inactive mutant protein xrn1-E178Q and rat1-E207Q respectively (Figure 2B). Moreover, the decay activity of both proteins on FAD-capped RNA was processive without discernable hydrolysis intermediates. In contrast, the *S. pombe* Rai1, which lacks exonuclease activity (25), only released the FAD cap without subsequent RNA degradation (Figure 2B). Collectively, these results suggest that both Xrn1 and Rat1 utilize the same catalytic active site for the hydrolysis of FAD capped RNA and for canonical 5’P RNA degradation.

**Figure 2.**
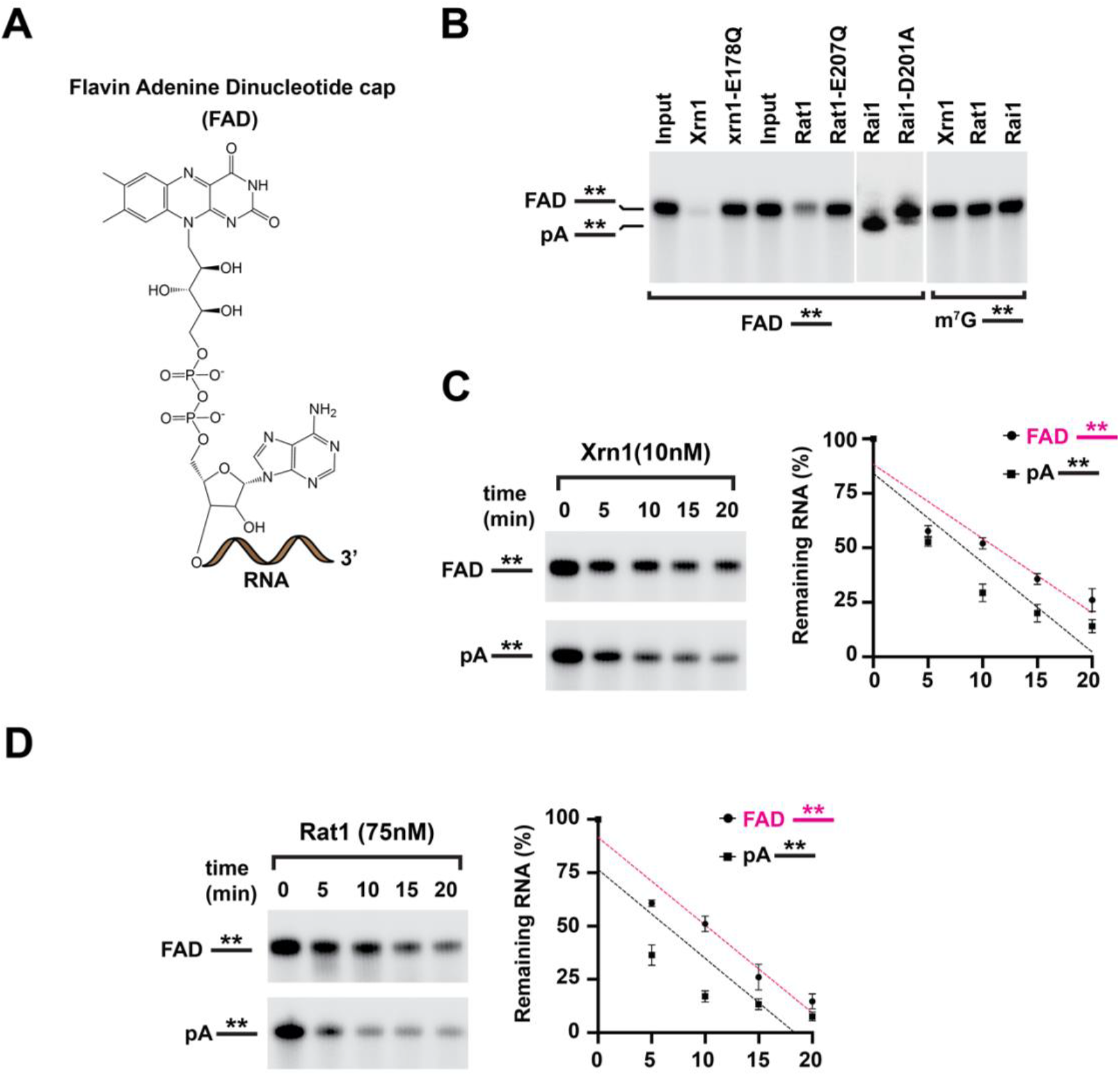
Xrn1 and Rat1 are deFADding enzymes. (A) Chemical structure of FAD-capped RNA. (B) Reaction products of *in vitro* deFADding assays with 50 nM recombinant Xrn1, WT, and catalytically inactive (*E178Q*) from *K. lactis or* 100 nM recombinant Rat1, WT or catalytically inactive (*E207Q*) from *S. pombe* or 25 nM Rai1, WT catalytically inactive (*D201A) (S. pombe*). Uniformly ^32^P-labeled monophosphate (pA–) or FAD-capped RNA (FAD–) where the line denotes the RNA were used as indicated. The asterisk represents the ^32^P-labeling within the body of the RNA. Reaction products were resolved on 15% 7M urea PAGE gels. Time-course decay analysis of uniformly ^32^P-labeled monophosphate or FAD-capped RNA with the indicated amount of Xrn1 (C) or Rat1 (D) protein are shown. Quantitation of RNA remaining is plotted from three independent experiments with ± SD denoted by error bars.

### Xrn1 possess higher deFADding activity compared to Rat1 *in vitro*

To assess the kinetics of deFADding activity relative to the well-established 5’ monophosphate (5’P) RNA degradation under the same assay conditions, we carried out time-course Xrn1 decay assays with 5’P- or FAD-capped RNAs. Intriguingly, the activity of Xrn1 on 5’P RNA was only ^~^1.5 fold more efficient than FAD-capped RNA at equivalent enzyme concentrations (Figure 2C). A comparison of Xrn1 hydrolysis of 5’P verses NAD-capped RNA revealed a ^~^20-fold slower decay of the latter relative to 5’P RNA (11) indicating a strong preference of Xrn1 to for FAD-capped RNA comparable to its propensity to degrade 5’P RNA.

To further delineate the differential preference of FAD-over NAD-capped RNA by Xrn1, its degradation activity was deciphered in the presence of competitors. Consistent with the decay assays, competition with free NAD or NAD3’p which would mimic the 5’ end of an NAD-capped RNA, were inefficient in their capacity to compete for the decay of either 5’P or FAD-capped RNAs (Supplementary Figure S2A). Minor competition was observed with self-competition when monitoring NAD-capped RNA. Although no competition was observed when using FAD or FAD3’P on 5’P RNA, inhibition of Xrn1 decay was evident on FAD-capped RNA and more prevalent on NAD-capped RNA (Supplementary Figure S2B). Moreover, the competition was more pronounced with the FAD-capped RNA mimic, FAD3’P compared to free FAD (Supplementary Figure S2B). Interestingly, inhibition was nevertheless detected when using higher concentration of FAD (Supplementary Figure S3). Collectively, the data demonstrate the deFADding activity of Xrn1 is more efficient than its deNADding activity and Xrn1 has a higher affinity to FAD-capped RNA compared to free FAD.

We next compared the activity of Rat1 on the hydrolysis of 5’P to that of FAD-capped RNA. As shown in Figure 2D, although higher concentrations of recombinant Rat1 are necessary to detect comparable decay, the efficiency of hydrolyzing 5’P RNA relative to FAD-capped RNA was ~2 fold, which was similar to that observed with Xrn1. Collectively, our results further expand the substrate RNAs regulated by Xrn1 and Rat1 beyond their well characterized 5’P substrate (26) and their recently described NAD-capped RNA hydrolysis (11).

### Loss of Xrn1 stabilizes FAD-capped RNA *in vitro* and *in vivo*

Having established the deFADding activity of recombinant Xrn1 *in vitro*, the impact of endogenous yeast Xrn1 on deFADding was assessed. *In vitro* decay of 5’ end RNA was carried out with extract derived from either wild-type (WT) or *xrn1Δ* cells using FAD-or m^7^G-capped RNAs immobilized and protected at the 3’-end with biotin-streptavidin. Exposure of the FAD-capped RNA to extract containing wild-type Xrn1 led to the decay of the RNA with an ~20-minute half-life, while a similar incubation with extract derived from *xrn1Δ* cells demonstrated a significant stabilization of the RNA (Figure 3A). Importantly, the decay rate observed with the *xrn1Δ* extract was comparable to that detected with m^7^G-capped RNA, which was predominantly refractory to Xrn1 deletion as expected. These data suggest endogenous Xrn1 possesses deFADding activity.

**Figure 3.**
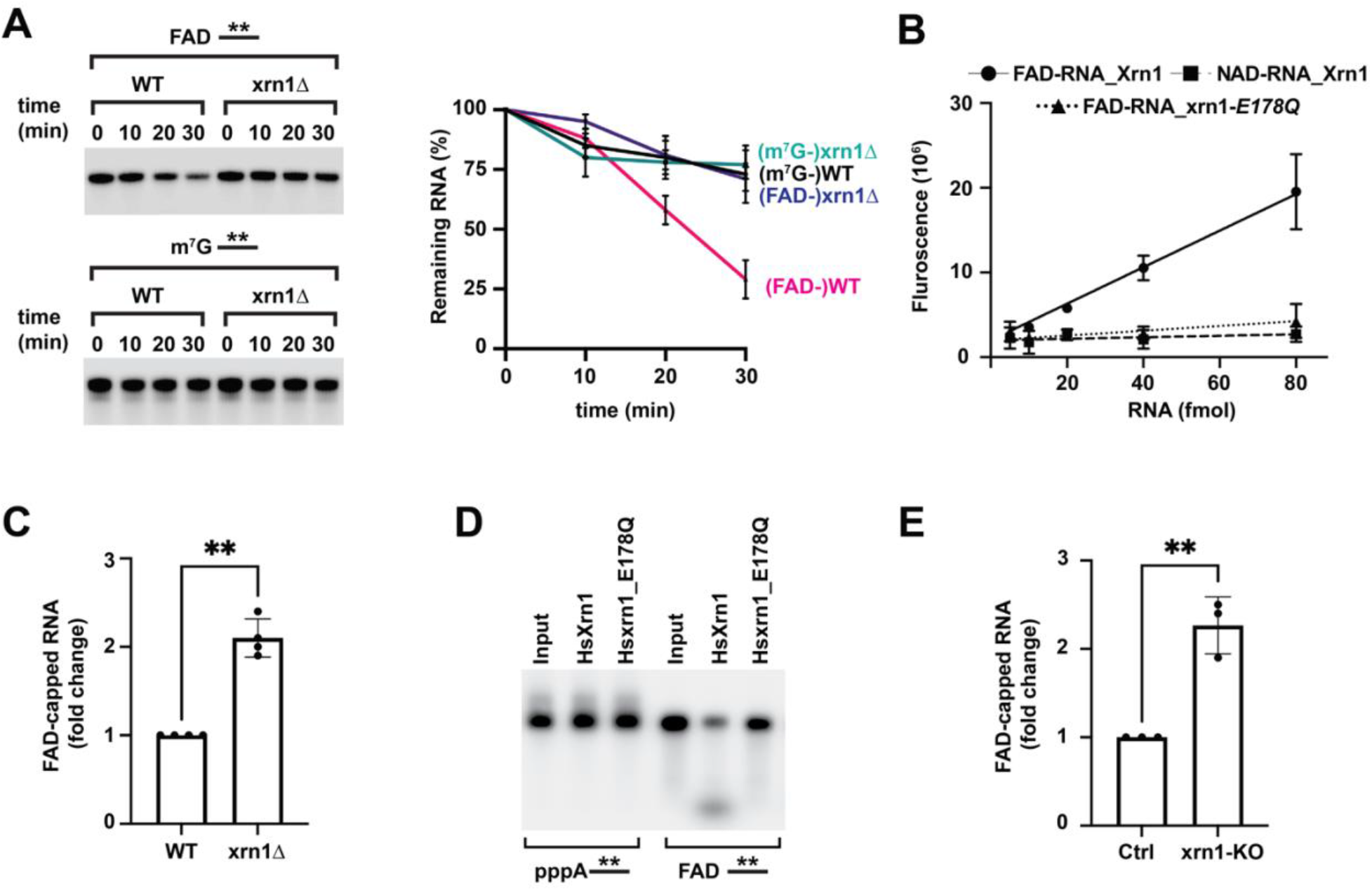
Xrn1 deFADs FAD-capped RNAs in both yeast and human cells. (A) Time-course decay analysis of uniformly ^32^P-labeled m^7^G- or FAD-capped RNA in the presence of cell extract prepared from WT or *xrn1Δ* strains. The remaining RNA was quantified and plotted from three independent experiments with ± SD denoted by error bars. (B) FAD-capQ assay using different amounts of *in vitro* transcribed FAD-capped RNA to demonstrate the release of intact FAD upon Xrn1 mediated deFADding. Catalytic-dead mutant of Xrn1, E178Q was used as a control for the Xrn1 reaction and NAD-capped RNA was used as a control for specific detection of FAD in the assay (C) Total RNAs from WT and *xrn1Δ* were subjected to the FAD-capQ assay to detect total levels of cellular FAD-capped RNA. Data represents average from three independent experiments. Error bars represent ± SD with unpaired *t*-test; * p < 0.05, ** p < 0.001, *** p < 0.0001 (D) Reaction products of *in vitro* deFADding assays with 100nM recombinant human Xrn1(Hs_Xrn1) and catalytically inactive (*E178Q*). (E) Small RNAs (<200 nts) from HEK293T Ctrl cells and xrn1-KO were subjected to the FAD-capQ assay to detect total level of cellular FAD-capped RNAs. Data represents average from three independent experiments. Error bars represent ± SD with unpaired *t*-test; * p < 0.05, ** p < 0.001, *** p < 0.0001.

To evaluate whether Xrn1 hydrolysed the FAD cap within the diphosphate moiety analogous to the class 2 Nudt12 protein (9), or removed the intact FAD, like the class 1 DXO/Rai1 family of proteins (7), FAD cap detection and quantitation (FAD-capQ) was used. FAD-capQ approach detects intact FAD caps released from the 5’-end of RNAs *en masse* (13). Hydrolysis of an increasing concentration of *in vitro* transcribed FAD-capped RNA with Xrn1 released a corresponding increase of FAD which was not observed in the reactions with catalytically dead Xrn1 mutant (Figure 3B). As expected, an FAD signal was also not observed when NAD-capped RNA was used. Next, we measured the contribution of Xrn1 deFADding on endogenous FAD-capped RNA. Consistent with a deFADding function for Xrn1 in cells, a statistically significant 2-fold higher level of total FAD cap was detected in the *xrn1Δ* strain relative to the WT strain (Figure 3C). These findings demonstrate Xrn1 is a class I deFADding enzyme that releases the intact FAD moiety, and this activity is also evident in cells.

The evolutionary conservation of Xrn1 deFADding beyond yeast was next assessed by testing recombinant and endogenous human Xrn1. As shown in Figure 3D, recombinant human Xrn1 hydrolyzed FAD-capped RNA, whereas the catalytically dead E178Q mutant did not. Similarly, the contribution of endogenous human Xrn1 to deFADding in cells was assessed using FAD capQ. Levels of FAD caps were found to be ~2 fold higher in human HEK293T cells with a CRISPR Cas9-directed disruption of the *Xrn1* gene (27)relative to the WT control cells (Figure 3E). We conclude the deFADding activity of Xrn1 is not restricted to yeast and at least extends to human Xrn1.

### Highly conserved Histidine 41 is dispensable for deFADding activity

Disruption of both deFADding and 5’P mediated exonuclease activities by the E178Q mutation in Xrn1 suggests both activities share catalytic residues for their enzymatic function. To assess whether the two activities could be uncoupled, we generated recombinant Xrn1 proteins with substitution mutations in several key residues within the catalytic center and in the 5’P binding pocket (Supplementary Figure S4A). No significant differences were detected with any of the proteins containing mutations in 4 critical residues in their ability to degrade FAD-capped or 5’P RNA *in vitro* (Supplementary Figure S4B). Unexpectedly, this also included a substitution of Histidine 41 to Alanine (H41A). The H41A mutation could distinguish between deNADding and exonuclease activities where deNADding activity was inhibited with minimal disruption of 5’P hydrolysis (11). Our findings reveal 5’P directed exonuclease activity and deFADding activities share common residues required for their catalytic function compared to the action of Xrn1 in deNADding.

### Bacterial 5’-3’ exoribonuclease RNase AM is a novel deFADding enzyme

The unexpected finding that both major eukaryotic 5’-3’ exoribonucleases Xrn1 and Rat1 possess deFADding and deNADding activity raised the possibility that metabolite cap decapping may be an intrinsic property of 5’ end exoribonuclease. To test this, we assessed the deFADding and deNADding activities of a recently characterized 5’-3’ exoribonuclease in *E. coli*, RNase AM (*yciV*) (14,15), which can hydrolyze short target RNAs up to 5 nucleotides with a 5’P. We were able to confirm the original observations with a 40 nucleotide RNA molecule and further demonstrate the exonuclease activity is processive (Figure 4A). Intriguingly, RNase AM also demonstrated robust deFADding activity, but unlike the two eukaryotic nucleases, RNase AM did not hydrolyze NAD-capped RNA (Figure 4A). Strikingly, RNase AM degraded both FAD-capped and 5’P RNAs at comparable rates *in vitro* (Figure 4B). Additionally, like Xrn1, RNaseAM degraded the FAD-capped RNA by removing and releasing the intact FAD as assessed by FAD-capQ (Figure 4C).

**Figure 4.**
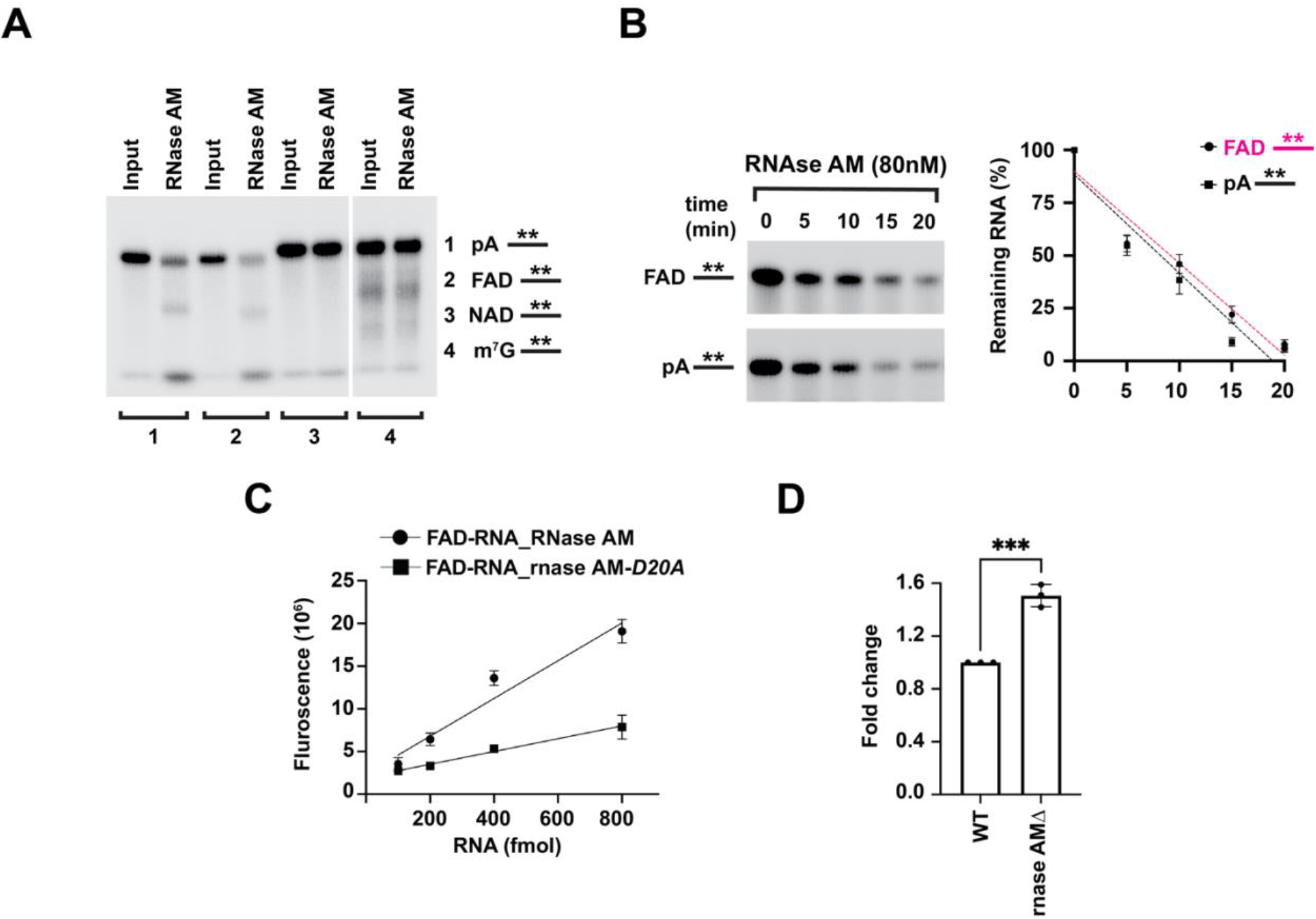
RNase AM is a novel deFADding enzyme in E. coli. (A) Reaction products of *in vitro* deFADding assays with 100nM recombinant RNase AM. Uniformly ^32^P-labeled monophosphate (pA–) or FAD-capped RNA (FAD–), NAD-capped (NAD–), and m^7^G-capped (m^7^G–). The asterisk represents the ^32^P-labeling within the body of the RNA. Reaction products were resolved on 15% 7M urea PAGE gels. (B) Time-course decay analysis of uniformly ^32^P-labeled monophosphate or FAD-capped RNA with the indicated amount of RNase AM protein are shown. Quantitation of RNA remaining is plotted from three independent experiments with ± SD denoted by error bars. (C) To demonstrate that RNase AM releases intact FAD upon deFADding like Xrn1 (in Fig 3), FAD-capQ assay using different amount of *in vitro* transcribed FAD-capped RNA was used. Catalytic-dead mutant of RNase AM, D20A was used as control for the reaction. (D) Total RNA from WT and *rnase AMΔ* were subjected to the FAD-capQ assay to detect total levels of cellular FAD-capped RNA. Data represents an average from three independent experiments. Error bars represent ± SD with unpaired *t*-test; * p < 0.05, ** p < 0.001, *** p < 0.0001.

The contribution of RNase AM on endogenous FAD-capped RNA was next determined. Total RNA from WT or *rnase AM* knockout *E. coli* strains were subjected to FAD-capQ and levels of released FAD detected. As shown in Figure 4D, a statistically significant 1.5-fold higher level of total FAD cap was detected in the *rnase AM* knock out strain relative to the WT strain. Our findings demonstrate that *E. coli* RNase AM can function as a deFADding enzyme in cells.

### Structural analysis of RNase AM

To elucidate the mechanism for RNase AM monophosphate hydrolysis and deFADding activity we attempted to determine the crystal structure of *E. coli* RNase AM in complex with FAD. RNase AM expressed in the presence of the iron chelator 2,2’-bipyridyl and MnSO_4_ was competent for binding FAD (as determined by thermal shift assay; data not shown). However, no crystals were obtained for RNase AM alone or in the presence of FAD. Good quality crystals were obtained of RNase AM incubated with a dsRNA, and a molecular replacement solution was obtained using the polymerase and histidinol phosphatase (PHP) domain from the *Chromobacterium violaceum* amidohydrolase, CV1693 (PDB: 2YB1) as the search model (21). No density for the RNase AM insertion domain was observed, indicating that the domain is disordered in this crystal. The final structure consists of residues 9-106 and 177-284 and is modeled with 3 manganese ions (Mna, Mnβ, and Mng) and a sulfate molecule in the active site. The refined atomic model contained no RNA and has excellent agreement with the X-ray diffraction data and the expected bond lengths, bond angles and other geometric parameters (Table 1).

RNase AM belongs to the PHP domain family of proteins which is composed of amidohydrolases/phosphoesterases and is present in some DNA polymerases. RNase AM has a distorted (β/α)_7_-barrel fold and a three metal enzyme active site composed of histidines and aspartates/glutamate residues (Figs. 5A and D). Mna and Mnβ, which are proposed to coordinate the water for nucleophilic attack are coordinated by residues His13, Glu72, and Asp255, and Glu72, His83 and His198 respectively (Fig. 5D). Mng is coordinated by residues Asp20, His45, and His257. The sulfate molecule coordinates all three metals and forms a salt bridge with Arg201, which may stabilize the hydrolysis reaction intermediate. Comparison to the CV1693 product complex containing AMP and P_i_ (21) shows that RNase AM has the same overall fold and active site architecture, and the RNase AM sulfate is in the same position as the phosphate product (Figs. 5C and D). Disruption of the active site by the D20A, E72A, and D255/H257A mutations resulted in disruption of 5’P and deFADding activities demonstrating the importance of the metals for these activities (Fig. 5G). Together these data indicate that RNase AM uses the same mechanism as CV1693 for hydrolysis.

**Figure 5.**
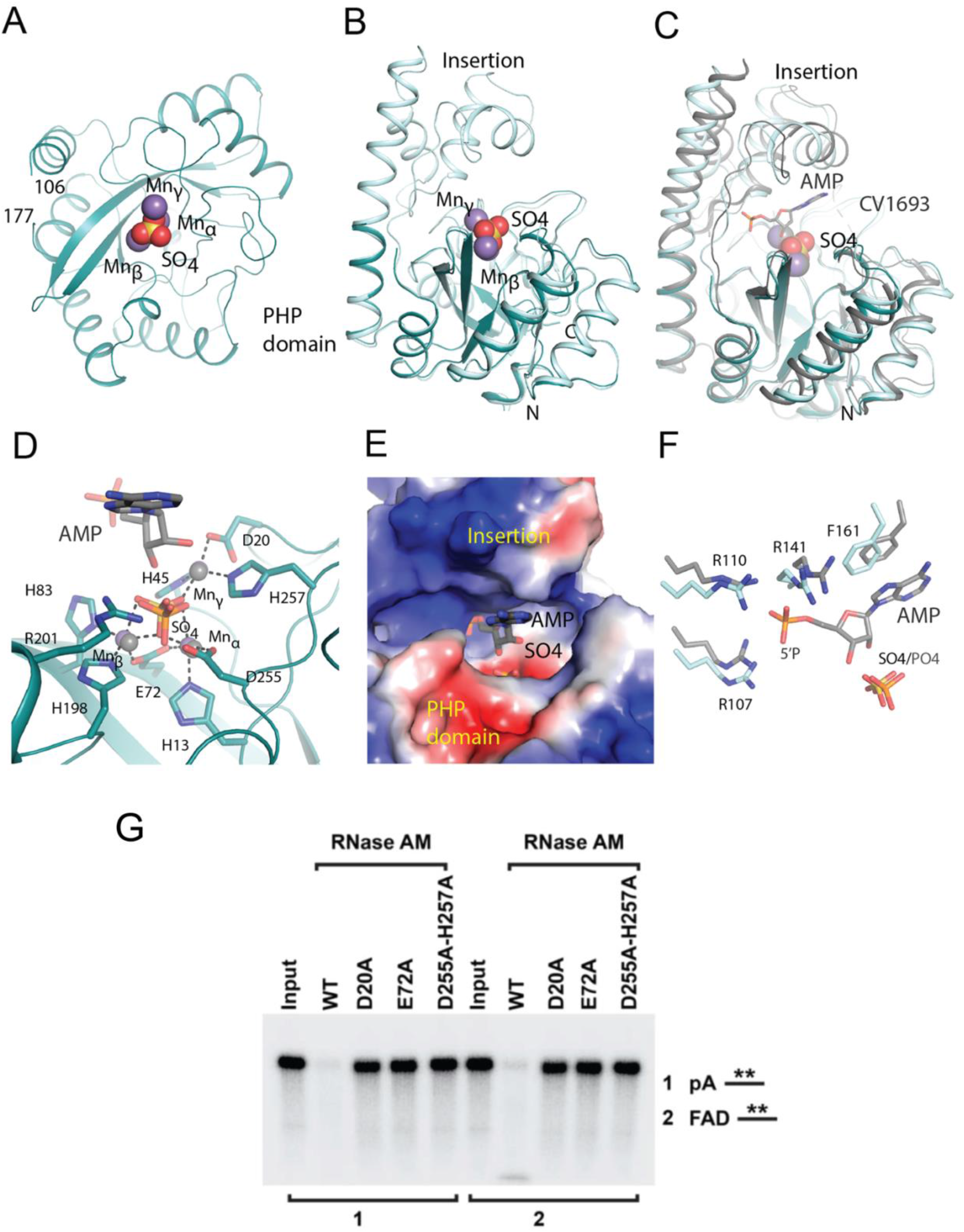
Structural and mutagenesis analysis of RNase AM activities. (A) Overall crystal structure of RNase AM PHP domain (teal) with manganese ions (purple) and sulfate (yellow and red) shown as spheres. (B) Overlay of RNase AM crystal structure (teal) with RNase AM from AlphaFold (light blue) showing the insertion domain. (C) Overlay of RNase AM with CV1693 (gray) with AMP from CV1693 shown as sticks. (D) The active site of RNase AM showing the coordination of the three manganese ions and sulfate to the active site residues (sticks). AMP and manganese (gray) and the phosphate (orange and red) from the overlay with CV1693 are shown. (E) Electrostatic potential mapped onto the surface of RNase AM showing the positively charged region where the 5’P from CV1693 AMP (gray) is buried. (F) Comparison of binding of AMP (gray) between RNase AM and CV1693 showing conservation of the residues that interact with AMP. (G) Reaction products of *in vitro* deFADding assays with 100 nM recombinant RNase AM and putative catalytically inactive point mutants. Uniformly ^32^P-labeled monophosphate (pA–) or FAD-capped RNA (FAD–) are indicated. The asterisk represents the ^32^P-labeling within the body of the RNA. Reaction products were resolved on 15% 7M urea PAGE gels

To gain further insight into the 5’P and deFADding activities of RNase AM we used the fulllength RNase AM model from AlphaFold (RNase AM_AF_). RNase AM_AF_ and the RNase AM PHP crystal structure align well with an RMSD of 0.58 Å and aligns with CV1693 with an RMSD of 1.71 Å (Figs. 5B and C). Comparison of RNase AM_AF_ with CV1693 shows that the 5’ phosphate of the CV1693 AMP (which represents the first nucleotide of 5’P RNA or the adenosine moiety from FAD) is buried in a positively charged cavity between the PHP and insertion domains comprised primarily of residues Arg107, Arg110, and Arg141 (Figs. 5E and F). Similar to CV1693, a stacking interaction can be made with the 5’ adenine base through Phe161. This mode of binding may explain 5’P activity of RNase AM; however the buried position of the 5’P indicates FAD cannot bind in the same manner, suggesting RNase AM utilizes distinct binding modes for 5’P and FAD RNA recognition.

## DISCUSSION

The 5’ cap of RNA is a critical determinant of the RNA fate. The N7-methyl-Guanosine (m^7^G) on the 5’end of mRNAs in eukaryotes is the most common and well-studied 5’ RNA cap (28). Recent highly sensitive chemical analysis of the 5’ ends of the RNA from prokaryotes, eukaryotes and pathogenic viruses has uncovered the presence of nucleotide metabolite caps (12). These can consist of adenosine derived metabolites– NAD, FAD, dpCoA and uracil derived sugars -UDP-GlcNAc and UDP-Glc. Other than the contribution of NAD caps to RNA stability, the function of the metabolite caps generally remains elusive. To begin exploring the function of FAD caps, we characterized proteins from the model organism, *S. cerevisiae* that can interact with the FAD cap. Surprisingly, the predominant FAD-cap interacting proteins were the 5’P dependent exoribonucleases that also possessed deFADding activity. Moreover, we demonstrated that the deFADding activity of 5’ end exoribonucleases was conserved throughout evolution and was also apparent in bacteria, in addition to yeast, and humans. RNase AM is so far the only 5’-3’ exoribonuclease known in *E. coli*, the enzymatic activity of which has been shown to be essential for the processing of bacterial ribosomal RNA (15).

We recently reported that both Xrn1 and Rat1 possess deNADding activity (11) in addition to their canonical monophosphate RNA hydrolysis activity (26,29). We further uncovered that that the deNADding activity of Xrn1 is essential for mitochondrial NAD capped RNA homeostasis (11). Remarkably, the deFADding activity of both Xrn1 and Rat1 was found to be more robust than their deNADding activity and as potent as their canonical monophosphate RNA hydrolysis activity. In contrast, the RNase AM exoribonuclease was more selective and exhibited deFADding, but not deNADding activity. Although we were able to uncouple the deNADding activity of Xrn1 from its monophosphate RNA decay by utilising a specific point mutation in the catalytic pocket, H41A, all our attempts to separate deFADding from monophosphate RNA hydrolysis for both Xrn1 and RNase AM were not fruitful. We screened several point mutations of residues that form the catalytic domain of both Xrn1 and RNase AM, along with the residues proximal to catalytic domain, but unfortunately could not untangle the enzymatic activities. Nevertheless, this implied that both activities are apparently mediated by identical residues and are inherent to 5’-3’ exoribonucleases.

Our analysis of the levels of FAD capped RNA isolated from both yeast and human Xrn1 knock out cells and RNase AM knock out cells assessed by FAD-capQ analysis validated that these 5’-3’ exoribonucleases can impact levels of total FAD-capped RNA in cells. These findings establish Xrn1 and RNase AM as novel deFADding enzymes adding to the existing list of enzymes that function in FAD cap decapping in cells including Rai1 (13), its mammalian homolog DXO, and Nudt16 (10). We also note that relative to NAD caps (12), FAD cap levels are significantly lower in yeast, human and bacterial cells as determined with FAD-capQ, which correlates well with the relative amount of free NAD and FAD in the cells (30,31).

At present it is difficult to decern the biological function of an FAD cap. Unlike a role for NAD caps in mitochondrial NAD homeostasis which was deciphered by uncoupling 5’P directed and NAD-cap directed decay (11), we were unable to separate the 5’P and deFADding activities (Supplementary Figure S4). Furthermore, although our initial objective was to find high affinity FAD cap-binding proteins that could be used to selectively identify FAD-capped RNAs, we surprisingly only identified nucleases rather than *bone a fide* high affinity cap binding proteins. The association of FAD caps with nucleases may suggest the cap functions as a mark for targeted decay analogous to the NAD cap in mammals. Further analysis of FAD cap function will require novel methodologies that can selectively isolate and identify cellular RNAs that harbor this noncanonical cap.

The structure of RNase AM provides important insights into the catalytic domain of this protein and its function as a 5’-3’ exonuclease. However, an RNase AM-FAD cocrystal structure will be necessary to discern the precise binding mode of this dinucleotide and the molecular basis of its deFADding activity. RNase AM is conserved in many pathogenic bacteria including *Klebsiella pneumoniae, Salmonella enterica, Vibrio cholerae, Yersinia pestis* (14) and the active site molecular details revealed in the present study could prove significant in small molecule targeting approaches.

## ACCESSION NUMBERS

Atomic coordinates and structure factors for the reported crystal structures have been deposited with the Protein Data bank under accession number XXXX. (to be provided at the proof stage).

## SUPPLEMENTARY DATA

Supplementary Data are available at NAR online.

## ACKNOWLEDGEMENT

We would like to thank Dr. Britt Glausinger (University California Berkley) for providing the HEK293T Xrn1 knockout cell line. We are grateful to Dr. Joanna Kowalska (University of Warsaw) for providing the FAD3’P dinucleotide. We also thank Dr. Haiyan Zheng (Rutgers University) for her extensive assistance in the analysis of mass spectrometry data.

