## Supplemental data for "Identification of a novel deFADding activity in 5’ to 3’ exoribonucleases"

### Supplementary Figures

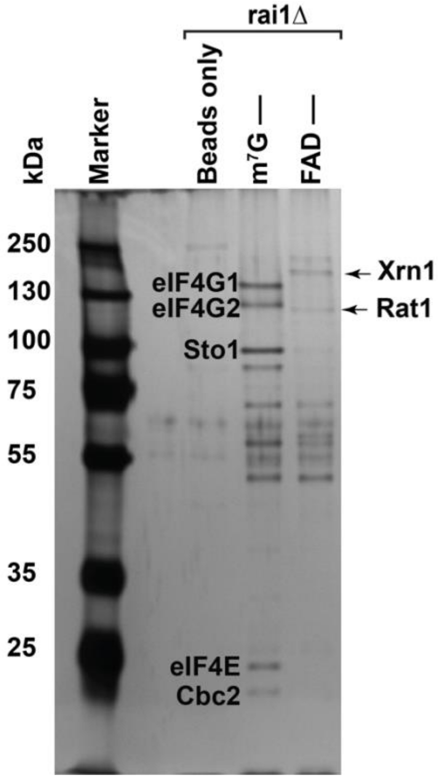


**Supplementary Figure S1. Characterization of proteins bound to the FAD cap in the absence of Rai1.** Protein derived from a strain lacking Rai1 (*rai1*Δ) were captured by FcRAP and eluates detected by Silver Staining following resolution on 10% SDS-PAGE gel. Affinity purification with 5′ $m^7G$ capped RNA ($m^7G$–) or FAD-capped RNA (FAD–) are shown. Streptavidin beads without RNA baits (beads only) was used a control.

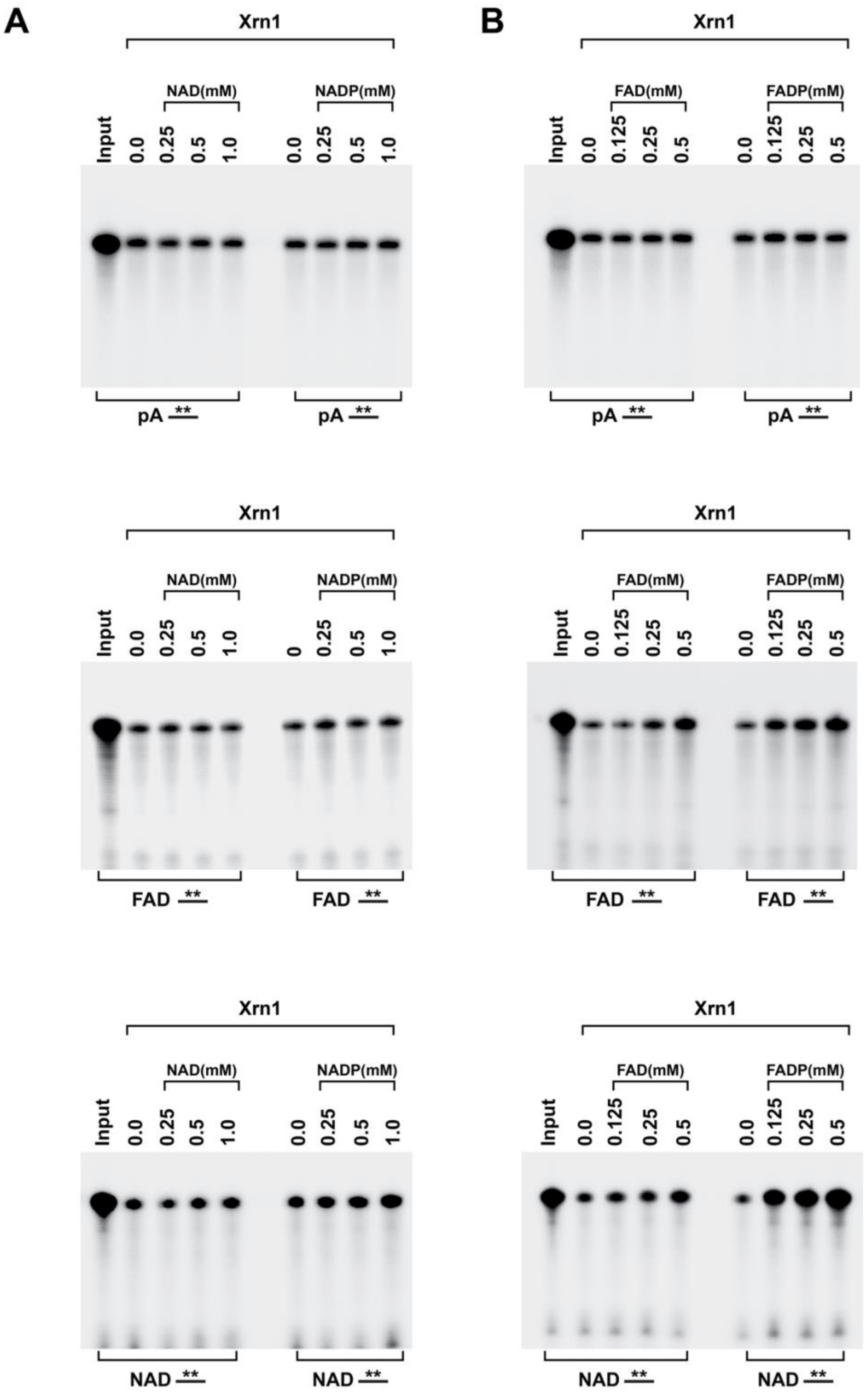


**Supplementary Figure S2. Xrn1 possess higher deFADding activity compared to its deNADding activity.** To define the differential preference of FAD- over NAD-capped RNA by Xrn1, its degradation activity was deciphered in the presence of competitors (free NAD or NAD3′p and free FAD and FAD3’p, which would mimic the 5′ end of an NAD-capped or FAD-capped RNA, respectively. 100nM recombinant Xrn1 from *K. lactis* was incubated with uniformly $^{32}P$-labeled 5′- monophosphate (pA–), NAD-capped or FAD-capped RNA in the presence or different amount of free NAD and NAD3'p **(A)**, FAD and FAD3'p **(B).** The products were resolved on 15% 7M urea PAGE gels

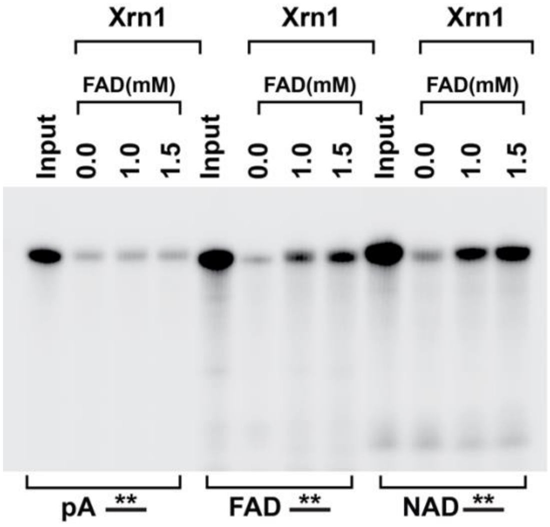


**Supplementary Figure S3. Higher concentration of free FAD inhibits deNADding and deFADding activity of Xrn1.** 100nM recombinant Xrn1 from *K. lactis* was incubated with uniformly $^{32}$P-labeled 5′- monophosphate (pA–), NAD-capped or FAD-capped RNA in the presence or different amount of free FAD**.** The products were resolved on 15% 7M urea PAGE gels.

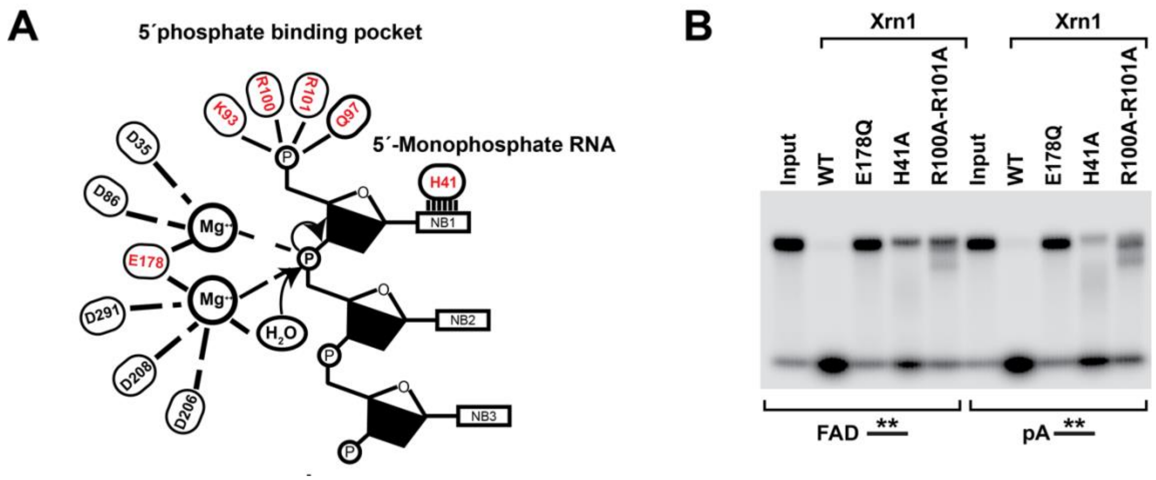


**Supplementary Figure S4. The xrn1-H41A point mutant is dispensable for FAD capped hydrolysis (A)** Model for monophosphate RNA hydrolysis (Jinek et al 2012). As shown recently (Sharma et al 2022), H41A abolishes the deNADding activity of Xrn1without affecting its 5’ monophosphate RNA hydrolysis. Surprisingly, as for monophosphate RNA, FAD capped RNA also remained unaffected upon mutating H41A. **(B)** 100nM recombinant WT or different point mutated Xrn1 from *K. lactis* was incubated with uniformly $^{32}$P-labeled 5′-monophosphate or FAD-capped RNA. The products were resolved on 15% 7M urea PAGE gels

### Supplementary Table S2. Oligonucleotides used in the present study

| NAME | SEQUENCE 5'->3' | PURPOSE |
|---|---|---|
| ϕ2.5-NAD/FAD-40 | CAGT**AATACGACTCACTATT**AGTTGGTGGTTGTTGTGTGTTTGTGGTTGGTTTGTTTGGC | In vitro transcription (T7 promoter sequence is highlighted) |
| PET-BACYV-A1 | TAAGAAGGAGATATACCATGAGCGACACGAATTATGCAGTGAT | For cloning YciV ORF in pET plamsids |
| PET-BACYV-A2 | GTGGTGGTGGTGGTGCTCGAGTAATTCCCTCTCTGTGGTGTTCTGCG | For cloning YciV ORF in pET plamsids |
| YCIV-D20A-A1 | CAGCTTCCGcTGGCTGCCTGACGCCAGAAG | For site directed mutagenesis (pET-rnase AM-D20A) |
| YCIV-D20A-A2 | CAGGCAGCCAgCGGAAGCTGTGGTATGGC | For site directed mutagenesis (pET-rnase AM-D20A) |
| YCIV-D255AH257A-A1 | GGATCTGcTTTTgcTCAGCCATGCCCGTGGATCG | For site directed mutagenesis (pET-rnase AM- D255AH257A) |
| YCIV-D255AH257A-A2 | GGCTGAgcAAAAgCAGATCCTTGTGATGCCCATAAATG | For site directed mutagenesis (pET-rnase AM- D255AH257A) |
| YCIV-E72A-A1 | CCCGGCGTGGcAATTTCCACGGTCTGGG | For site directed mutagenesis (pET-rnase AM- E72A) |
| YCIV-E72A-A2 | CGTGGAAATTgCCACGCCGGGAATAAGATTC | For site directed mutagenesis (pET-rnase AM- E72A) |

### Supplementary Table S3. Reagents used in the present study

| REAGENT or RESOURCE | SOURCE | IDENTIFIER |
|---|---|---|
| **Yeast/E.coli Strains and Human Cell lines** | | |
| BY4741 (MATa; his3Δ1; leu2Δ0; met15Δ0; ura3Δ0) | Dharmacon | YSC1048 |
| xrn1Δ (BY4741; MATa; his3Δ1; leu2Δ0; met15Δ0; ura3Δ0; Xrn1: KanMX | This study | NA |
| *E. coli* K-12 BW25113 | Horizon Discovery | OEC5042 |
| yciv/rnase AM-KO | Horizon Discovery | OEC4987-200826696 |
| HEK293T | Gilbertson et al (2018) (1) | NA |
| Xrn1-KO | Gilbertson et al (2018) (1) | NA |
| **Plasmids** | | |
| pET-KlXrn1 (A derivative pET26a(+) plasmid carrying KlXrn1(1,1245)-6xHis fusion protein) | Chang et al(2011) (2) | NA |
| pET-klxrn1-H41A (A derivative pET-KlXrn1 plasmid carrying mutant klxrn1-H41A-6xHis fusion protein) | Sharma et al (2022) (3) | |
| pET-RNAse AM (A derivative pET26a(+) plasmid carrying YciV(ORF)-6xHis fusion protein | This study | NA |
| pET-rnase AM-D20A (A derivative pET26a(+) plasmid carrying mut carrying mutant yciV-D20A -6xHis fusion protein | This study | NA |
| pET- rnase AM-E72A (A derivative pET26a(+) plasmid carrying mutant yciV-E72A -6xHis fusion protein | This study | NA |
| pET- rnase AM-D255A-H257A (A derivative pET26a(+) plasmid carrying mutant yciV- D255A-H257A -6xHis fusion protein | | |
| pET-klxrn1-K93A (A derivative pET-KlXrn1 plasmid carrying mutant klxrn1-K93A-6xHis fusion protein) | Sharma et al (2022) (3) | NA |
| pET-klxrn1-Q97A (A derivative pET-KlXrn1 plasmid carrying mutant klxrn1-K97A-6xHis fusion protein) | Sharma et al (2022) (3) | NA |
| pET-HsXrn1 (A derivative pET26a(+) plasmid carrying HsXrn1(1,1189)-6xHis fusion protein) | Sharma et al (2022) (3) | NA |
| pET-HsXrn1-E178Q (A derivative pET26a(+) plasmid carrying mutant HsXrn1-E178Q(1,1189)-6xHis fusion protein) | Sharma et al (2022) (3) | NA |
| pET-klxrn1-R100/101A (A derivative pET-KlXrn1 plasmid carrying mutant klxrn1-R100A, R101A-6xHis fusion protein) | Sharma et al (2022) (3) | NA |
